# Proteasome Inhibition Reduces NBR1 Protein Levels in mATG8-Deficient Cells Independently of Lysosomal Degradation and RAB27A

**DOI:** 10.1101/2025.07.15.664987

**Authors:** Cristóbal Cerda-Troncoso, Eloísa Arias-Muñoz, Gabriela Vargas, Belén Gaete-Ramírez, Karina Cereceda, Viviana A. Cavieres, Patricia V. Burgos

## Abstract

Autophagy is a lysosome-dependent degradation process that involves autophagosome formation, typically mediated by mammalian ATG8 proteins (mATG8s). Autophagosomes can still form in their absence, suggesting alternative mechanisms. NBR1, a selective autophagy receptor, has been shown to compensate for autophagy defects. Here, we examined NBR1 regulation under proteotoxic stress in mATG8s-deficient HeLa cells. NBR1 levels were elevated in mATG8s knockout (KO) cells under basal conditions but decreased significantly after treatment with the proteasome inhibitor MG132. This reduction was not prevented by lysosomal inhibition with BafA1, indicating a non-lysosomal mechanism. Silencing of RAB27A reduced basal NBR1 levels, and the effects of MG132 were no longer observed, likely due to already diminished NBR1. Remaining NBR1 localized to puncta positive for ubiquitin and the ESCRT-0 component HRS, suggesting involvement of a ubiquitin-dependent endosomal pathway. Overall, our results suggest that under proteotoxic stress and impaired autophagy, cells activae alternative routes, potentially involving unconventional secretion, to regulate NBR1 levels.

**Graphical abstract:** 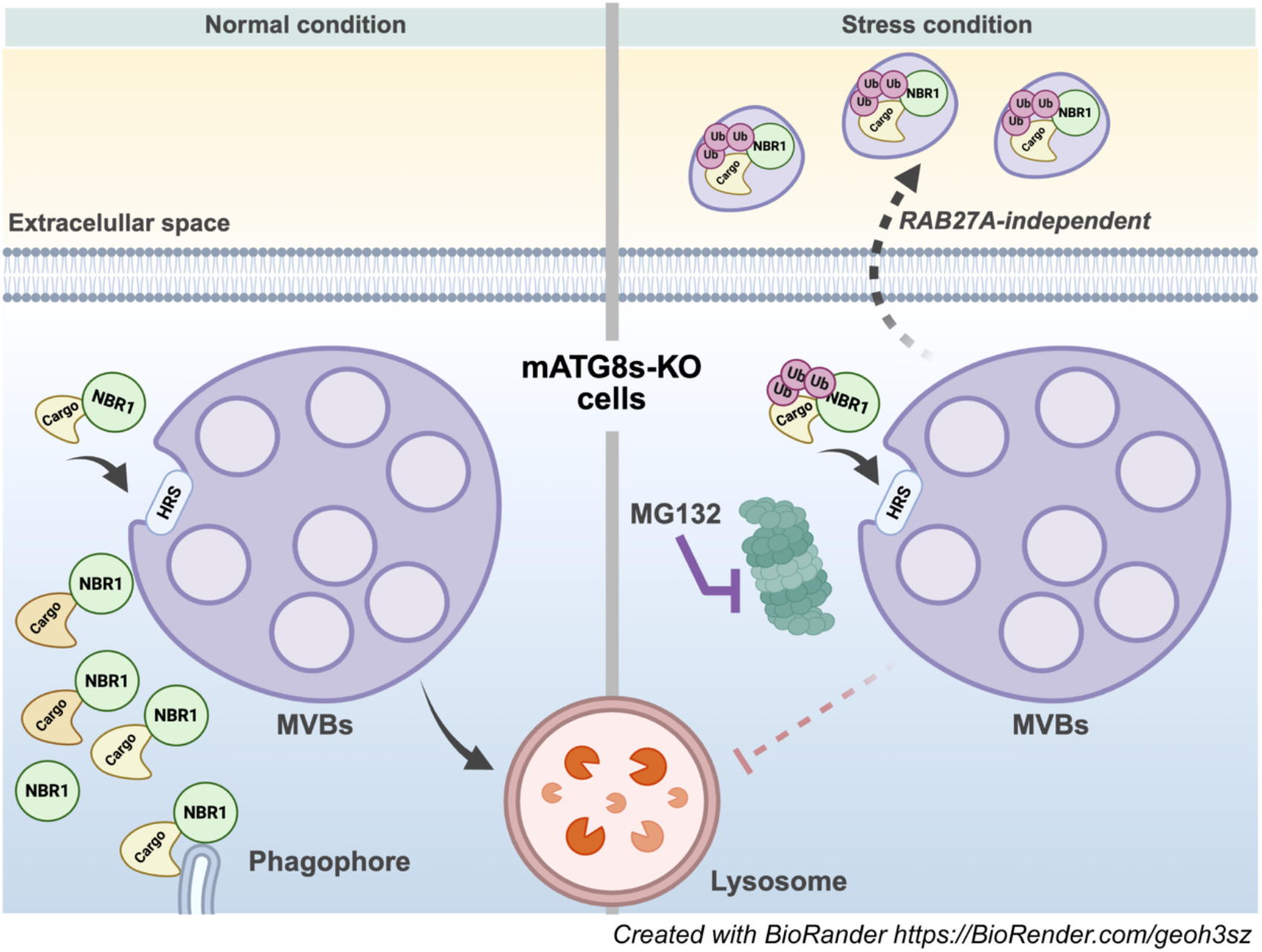

Created with BioRender https://BioRender.com/geoh3sz

In the absence of mATG8s, NBR1 may be recruited to multivesicular bodies (MVBs) via HRS, for later degradation in lysosomes (left panel). Under proteasomal inhibition (right panel), NBR1 associated with MVBs (with ubiquitinated cargos) is downregulated, independent of lysosomes. Possibly, this downregulation is due to a secretion process, through a RAB27A-independent mechanism.

## Introduction

(Macro)Autophagy is a catabolic process that delivers cytoplasmic components to the lysosome for degradation through double-membrane autophagosomes. Its biogenesis requires the coordinated action of autophagy-related (ATG) proteins. In yeast, ATG8 is essential for autophagosome formation (1), while in mammals, this function is fulfilled by ATG8 orthologs (mATG8s), including LC3 and GABARAP protein families. Notably, unlike yeast, mammals can form autophagosomes even in the absence of mATG8s (2). Selective cargo recognition is mediated by autophagy receptors such as NBR1 and SQSTM1/p62 (3). In the absence of mATG8s, cargo can instead be secreted via autophagy-dependent secretion (4). Notably, NBR1 compensates for the loss of ATG7, an essential protein involved in mATG8s lipidation by promoting autophagosome formation through an alternative mechanism (5).

Given the central role of NBR1 in coordinating selective autophagy and potential compensatory pathways, we explored how NBR1 behaves under conditions of proteasomal inhibition in the absence of mATG8s. For this, HeLa cells, both WT and mATG8s KO, were compared, revealing that mATG8s KO cells displayed elevated levels of NBR1 relative to WT. Surprisingly, proteasomal inhibition with MG132 led to a significant decrease in NBR1 levels in mATG8s KO cells. Inhibiting lysosomal degradation using BafA1 failed to prevent this reduction, suggesting a non-lysosomal mechanism. Moreover, silencing of RAB27A, a GTPase involved in endo-lysosomal vesicle trafficking and secretion, reduced basal NBR1 levels and abolished its further decrease by MG132, likely due to already diminished levels rather than a requirement for RAB27A. Residual NBR1 localized in punctate structures positive for ubiquitin and the ESCRT-0 component HRS, suggesting a ubiquitin-dependent endosomal mechanism. Altogether, these findings suggest that under proteotoxic stress and impaired autophagy, cells engage alternative pathways, potentially involving unconventional secretion of NBR1.

## Methods

### Reagents and Antibodies

MG132 (cat#47479), Bafilomycin A1 (BafA1, cat#B1793), anti-NBR1 (cat#sc-130380, NBP171703), anti-EEA1 (cat#61045), anti-CD63 (cat#ab8219), anti-Cathepsin D (cat#AF1014), anti-HRS (cat#ab15539), anti-ubiquitin (cat#AUB01), and anti-RAB27A (cat#168013) were used.

### Cell Culture, Treatments, and Western Blot

HeLa WT and mATG8s KO cells (courtesy of Dr. Maho Hamasaki, Osaka University), and HEK293T cells (cat#632180), were cultured in DMEM High Glucose (cat#12800-017) supplemented with 10% heat-inactivated FBS (cat#sv30160.03), 100 U/mL penicillin, and 100 μg/mL streptomycin (cat#15070-063), at 37°C and 5% CO_2_. Cells were seeded after trypsinization with 0.05% trypsin and 1 mM EDTA (cat#25300-54).

For treatments, cells were incubated with 10 μM MG132 for 8 h or 100 nM BafA1 for 4 h. DMSO was used as vehicle control.

Cells were washed with cold PBS (cat#46-013) and lysed in RIPA buffer [50 mM Tris-HCl, 150 mM NaCl, 5 mM EDTA, 1% NP-40, 1% sodium deoxycholate, 0.1% SDS, pH 7.4] supplemented with protease inhibitors (cat#P8340). Lysates were sonicated on ice (3 pulses, 2–3 s, 40 mA) and processed for SDS-PAGE, Western blotting, and membrane detection following previously described protocols (6).

### Immunofluorescence and Confocal Microscopy

Cells grown on coverslips were fixed, permeabilized, and stained as previously described (6). Images were acquired using a Leica TCS SP8 confocal microscope (6).

### Lentiviral Transduction and Stable Knockdown

RAB27A knockdown was achieved by lentiviral transduction with shRNA as previously described (6). mATG8s KO cells were selected with 6 μg/mL puromycin (cat#P8833). Knockdown efficiency was confirmed by Western blot.

### Data Analysis

Densitometry and image quantification were performed using FIJI (ImageJ), Excel 2022, and Prism 10.0 (macOS). Statistical significance was determined using a one-tailed t-test.

## Results

### Proteasomal inhibition with MG132 reduces NBR1 levels in HeLa cells lacking mATG8s proteins, independently of lysosomal degradation

The absence of mATG8s in HeLa cells perturbs autophagosome trafficking, alters their morphology, and slows down their biogenesis rate (2), ultimately leading to the accumulation of selective autophagy receptors. Consistent with this, we confirmed the accumulation of NBR1 in mATG8s-KO HeLa cells compared to wild-type (WT) by Western blot analysis (Fig. 1A, lanes 1 and 3; Fig. 1B). Considering that in mATG8s-KO cells, cells may rely on the proteasomal system to maintain proteostasis, where NBR1 can promote substrate delivery to the proteasome, we explore whether proteasome inhibition increases NBR1 levels. Contrary to our expectations, treatment with 10 μM MG132 for 8 h led to a significant decrease in NBR1 protein levels in mATG8s-KO cells, as shown by Western blot (Fig. 1A line 3 and line 4; Fig. 1B) and immunofluorescence (Fig. 1C and 1D), a phenotype that was not observed in WT cells. Proteasome inhibition is known to activate lysosomal biogenesis and autophagosome degradation (7). We tested whether the reduction in NBR1 levels was due to lysosomal degradation. Cells were treated with 10 μM MG132 for 8 h, with 100 nM BafA1 added during the final 2 h to inhibit lysosomal activity. Lysosomal inhibition did not restore NBR1 levels in mATG8s-KO cells (Fig. 1E and 1F), indicating that the MG132-induced decrease in NBR1 is not mediated by lysosomal degradation.

**Figure 1.**
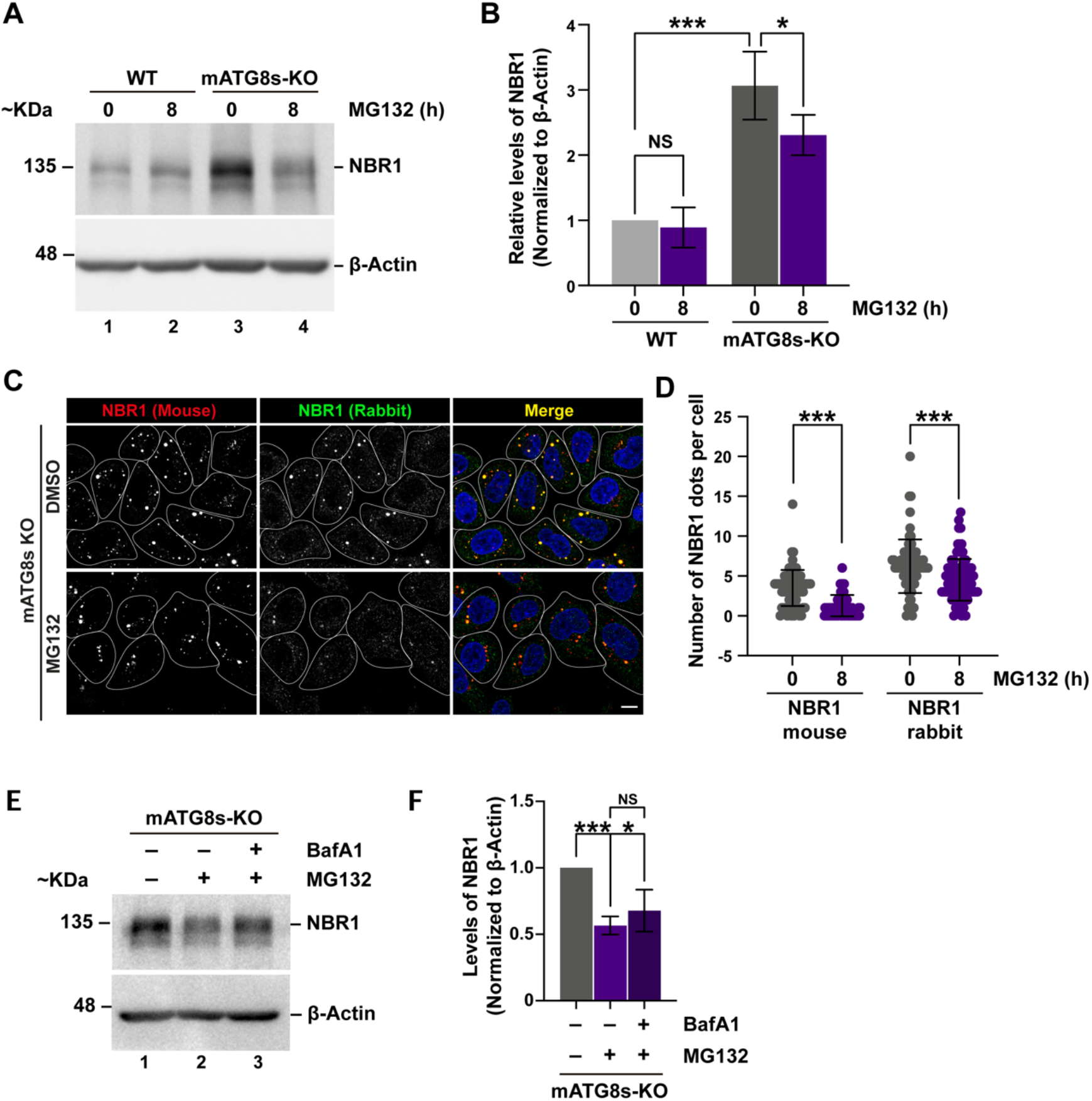
Catalytic proteasome inhibition reduces NBR1 levels in mATG8-deficient cells. (A) Western blot of endogenous NBR1 and β-actin in HeLa WT and HeLa mATG8s KO cells treated with 10 μM MG132 for 0 or 8 h. Cells were lysed with RIPA buffer. (B) Densitometric quantification of NRB1 levels shown in (A). NS: No-Significative **P<0.01, ***P<0,001; unpaired-parametric t-Test. (C) Immunofluorescence analysis of NBR1 distribution in HeLa WT and HeLa mATG8s-KO cells treated with 10 μM MG132 for 0 or 8 h; Scale bar 10 μm. (D) Quantification of NBR1 positive puncta per cell from images in (C); ***P<0,001; unpaired-parametric t-Test. (E) Western blot of endogenous NBR1 and β-actin in HeLa mATG8s KO cells treated with 10 μM MG132 for 0 or 8 h, with or without 100 nM BafA1 added during the final 2 h. (F) Densitometric quantification of NBR1 levels from (E); NS: No-Significative, *P<0.05, ***P<0.001; unpaired-parametric t-Test.

### Residual NBR1 levels in response to catalytic proteasome inhibition co-localize with ESCRT-0 component HRS and Ubiquitin aggregates

To gain insight into the trafficking pathways involved, we next assessed the subcellular distribution of NBR1 following MG132 treatment. Given that NBR1 can be incorporated into endosomes under basal conditions in yeast (8), we evaluated its co-localization with markers of the endolysosomal pathway in mATG8s-KO cells treated with MG132, including early endosomes (EEA1), the ESCRT-0 component HRS (involved in endosomal sorting (9)), late endosomes (CD63), and lysosomes (Cathepsin D, CathD). Confocal microscopy analysis revealed a significant co-localization of NRB1 with HRS, compared to the other markers (Fig. 2A and 2B), predominantly within large circular structures (Fig. 2A, arrowheads). Since NBR1 recognizes ubiquitinated cargos for degradation, we next examined the co-localization of NBR1 and HRS with ubiquitin following proteasome inhibition in mATG8s-KO cells. Confocal analysis showed that both NBR1 and HRS co-localize with ubiquitin after MG132 treatment (Fig. 2C and 2D), within similar structures.

**Figure 2.**
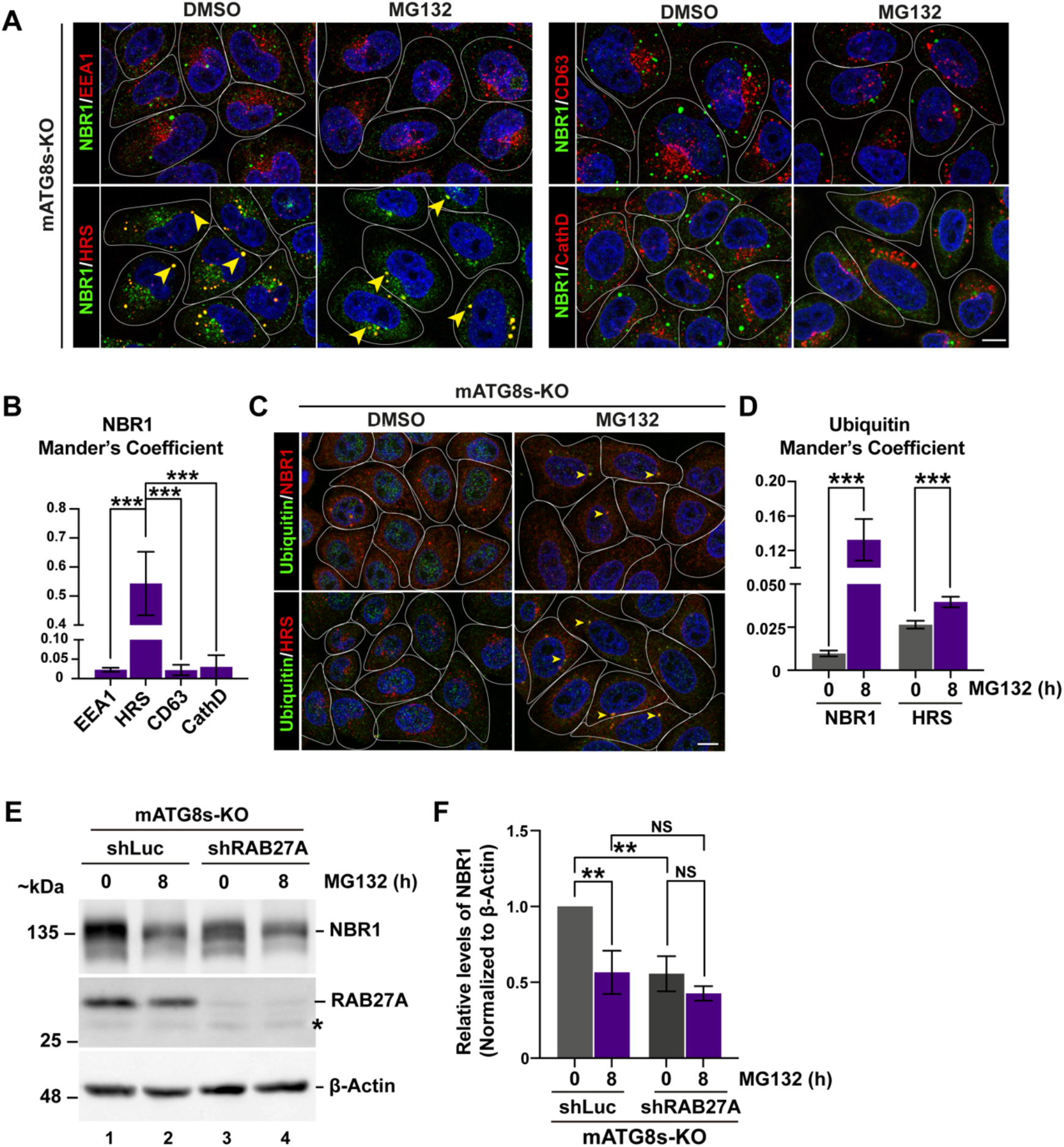
Residual NBR1 localizes to HRS and ubiquitin structures following catalytic proteasome inhibition. (A) Confocal analysis of NBR1 with endolysosomal markers in mATG8s KO cells treated with 10 μM MG132 for 0 or 8 h. Markers include EEA1 (early endosomes), HRS (MVBs), CD63 (secretory MVBs), and CathD (lysosomes); arrowheads indicate areas of co-localization. (B) Representative quantification of NBR1 co-localization with HRS from panel (A). (C) Mander’s coefficient quantification of NBR1 co-localization with the indicated endolysosomal markers after MG132 treatment; ***P<0.001; unpaired-parametric t-Test. (D) Confocal analysis of NBR1 or HRS co-localization with ubiquitin after 0 or 8 h of 10 μM MG132 treatment in mATG8s KO cells; arrowheads show co-localization. (D) Mander’s coefficient quantification of ubiquitin co-localization with NBR1 or HRS following MG132 treatment as shown in (C); ***P<0.001; unpaired-parametric t-Test. (F) Western blot analysis of NBR1 and β-actin protein levels in mATG8s KO either shLuc and shRAB27A cells, treated with 10 μM MG132 for 0 or 8 h. (G) Densitometric quantification of Western blot shown in (F); NS: No-Significative **P<0.01; unpaired-parametric t-Test.

Proteasomal inhibition with MG132 reduces NBR1 levels in mATG8-KO cells, via a RAB27A-independent mechanism.

Given the observed association between HRS and NBR1 after proteasome inhibition, and that HRS mediates earlier steps in intraluminal vesicle formation at endosomes, later released as exosomes via MVB-plasma membrane fusion regulated by the GTPase RAB27A (9), we investigated whether NBR1 reduction could be mediated by RAB27A-dependent secretion. To test this, we generated stable RAB27A knockdown in mATG8s-KO cells via lentiviral transduction (shRAB27A), using cells expressing a shRNA against luciferase as a control (shLuc). Western blot analysis revealed reduced basal NBR1 levels in shRAB27A compared to shLuc cells (Fig. 2E, lanes 1 and 3; Fig. 2F). We next assessed the effect of proteasome inhibition in these lines. As expected, MG132 treatment reduced NBR1 levels in shLuc cells (Fig. 2E, lanes 1 and 2; Fig. 2F). In contrast, MG132 treatment did not further decrease NBR1 levels in shRab27A cells (Fig. 2E, lanes 3 and 4; Fig. 2F), suggesting that the reduction of NBR1 upon proteasomal inhibition occurs through a RAB27A-independent mechanism.

## Discussion

Here, we show that NBR1 protein levels, elevated in mATG8s-KO cells, decrease under two distinct stress conditions: proteasomal inhibition and RAB27A silencing. The remaining NBR1 signal localizes to dot-like structures that co-localize with the ESCRT-0 component HRS and ubiquitin, suggesting involvement of the ubiquitin-dependent endosomal pathway. Consistent with this, lysosomal inhibition using BafA1 did not restore NBR1 levels, and no intracellular accumulation was observed, supporting the notion that its reduction is not due to lysosomal degradation. RAB27A silencing in mATG8s-KO cells also reduced NBR1 levels, suggesting that in the absence of this GTPase, NBR1 may be rerouted toward degradation or secretion via yet unidentified mechanisms. Although we did not directly assess secretion, our findings align with the possibility that NBR1 is released via a non-canonical pathway under cellular stress. The absence of MG132-induced NBR1 reduction in RAB27A-silenced cells likely reflects already diminished basal levels due to RAB27A depletion, rather than indicating its requirement in the MG132 response. Thus, NBR1 downregulation upon proteasome inhibition likely proceeds through a RAB27A-independent mechanism.

These results suggest that when canonical autophagy is compromised, cells become more vulnerable to additional stressors such as proteasome inhibition or vesicular trafficking disruption. In compensation, they may activate alternative pathways involving selective autophagy receptors that bind ubiquitinated cargo, with HRS potentially contributing. HRS has been implicated in autophagy-dependent secretory autophagy, typically requiring LC3 (10). However, our data point to an LC3-independent mechanism, consistent with the absence of mATG8s, where NBR1 may be redirected into ubiquitin-enriched vesicles. Whether these vesicles are ultimately secreted, and the nature of the underlying pathway, remain to be determined.

Given that aging and chronic proteotoxic stress impair canonical autophagy, identifying alternative routes for protein handling may offer new insights into cellular resilience and protein quality control.

## Acknowledgments

We kindly thank to Drs. Manuel Varas-Godoy (Universidad San Sebastián) and Matías Ostrowski (Universidad de Buenos Aires) for providing the plasmids used in lentiviral particle production. We are also grateful to Dr. Nicolás Albornoz for his contribution to the generation of raw ∼∼∼+∼data not included in this work.

## Author contributions

Conceptualization, **C.C-T., V.A.C** and **P.V.B**.; Methodology, **C.C.T., A.E-M., B.G-R., K.C**., and **V.A.C**; Formal analysis, **C.C-T., A.E-M., V.A.C**., and **P.V.B**.; Resources, **C.C-T., V.A.C**., and **P.V.B**.; writing—original draft preparation, **C.C-T., V.A.C**., and **P.V.B**. writing— review and editing, **C.C-T., V.A.C**., and **P.V.B**.; funding acquisition, **V.A.C**., and **P.V.B**.

